# A small excitation window allows long-duration single-molecule imaging, with reduced background autofluorescence, in *C. elegans* neurons

**DOI:** 10.1101/2022.12.20.521330

**Authors:** Aniruddha Mitra, Elizaveta Loseva, Guus H. Haasnoot, Erwin J.G. Peterman

## Abstract

Single-particle imaging using laser-illuminated widefield epi-fluorescence microscopy is a powerful tool to investigate molecular processes *in vivo*. Performing high-quality single-molecule imaging in such biological systems, however, remains a challenge due to difficulties in controlling the number of fluorescing molecules, photobleaching, and the autofluorescence background. Here, we show that by exciting only a small, 5-15 µm wide region in chemosensory neurons in live *C. elegans*, we can significantly improve the duration and quality of single-molecule imaging. Small-window illumination microscopy (SWIM) allows long-duration single-particle imaging since fluorescently labelled proteins are only excited upon entering the small excited area, limiting their photobleaching. Remarkably, we also find that using a small excitation window significantly improves the signal-to-background ratio of individual particles. With the help of theoretical calculations, we explain that the improved signal-to-background ratio is due to reduced background, mostly caused by out-of-focus autofluorescence. We demonstrate the potential of this approach by studying the dendritic transport of a ciliary calcium channel protein, OCR-2, in the chemosensory neurons of *C. elegans*. We reveal that OCR-2-associated vesicles are continuously transported back and forth along the length of the dendrite and can switch between directed and diffusive states. Furthermore, we perform single-particle tracking of OCR-2-associated vesicles to quantitatively characterize the transport dynamics. SWIM can be readily applied to other *in vivo* systems where intracellular transport or cytoskeletal dynamics occur in elongated protrusions, such as axons, dendrites, cilia, microvilli and extensions of fibroblasts.

## Introduction

Living organisms and cells are complex and heterogeneous systems where myriads of processes take place at the same time. The same protein can have different dynamics and functionality, depending on its subcellular localization, conformational state, chemical modification, and presence of interacting partners. Fluorescence microscopy is a great tool for investigating protein dynamics in real-time in their native context [1, 2]. Dynamics of clearly defined structures, such as microtubule bundles, mitotic chromosomes and nuclei can be revealed by means of fluorescently labelling the whole pool of a protein of interest and imaging the bulk fluorescence signal. This generally provides an averaged image of all of these protein molecules, which might keep the stochastic behaviour of individual protein molecules hidden, limiting important insights in the molecular basis of processes of interest. Single-molecule fluorescence microscopy and analysis techniques allow overcoming these limitations and provide spatial information with sub-diffraction limit precision [3–6].

The quality of single-molecule information depends on the number and photostability of emitting fluorophores, signal-to-background (SBR) and signal-to-noise (SNR) ratios, and rate of image acquisition. All these parameters are intertwined and require optimization for each particular experiment. A sparse density of fluorescing proteins, which is required for tracking and analysing them individually, can be achieved by labelling the proteins with photoactivatable or photoconvertible fluorescent markers and activating only a small fraction of them [3, 7–9]. Alternatively, one can start with many fluorescently-labelled proteins and photobleach them until only a few of them remain fluorescent in the region of interest [10]. The SBR ratio can be enhanced by using bright and photostable fluorescent proteins and by reducing the background (BG) that is caused by out-of-focus fluorescence and autofluorescence. Autofluorescence is due to other fluorescing molecules in the sample than the label and is inherent to most biological samples. The autofluorescence spectrum depends on the particular type of cell or tissue and its metabolic state. Generally, it is excited by wavelengths in the ultraviolet and blue range and emitted over a wide range of wavelengths in the visible spectrum [11–13]. Imaging conditions have to be optimized in order to collect the maximum fluorescence signal of the labels while minimizing autofluorescence. Out-of-focus fluorescence can be eliminated by exciting only a thin layer of the sample, close to the focal plane of the objective – a principle underlying techniques such as Total Internal Reflection Fluorescence (TIRF) microscopy, Highly Inclined and Laminated Optical sheet (HILO) microscopy, Light Sheet Fluorescence Microscopy (LSFM), which combine video-rate acquisition speeds, with improved SBR [14–18].

In earlier studies, we have shown that it is possible to image and track single molecules within cilia and dendrites of chemosensory neurons in the multicellular organism *C. elegans* (illustrated in Figure 1A) [19–23]. This could be done using laser-illuminated widefield epi-fluorescence microscopy (which is very sensitive to out-of-focus background fluorescence), since the transparent worms have a limited number of ciliated neurons. Furthermore, the dendrites and cilia are thin (100-500 nm) and contain a specific subset of proteins not expressed in other cells. To image ciliary proteins, we endogenously label proteins with fluorescent proteins such as eGFP and mCherry. To reach the single-molecule level we use photobleaching to reduce the number of fluorescing proteins, such that only a few molecules are visible in the imaged area at any given time. This strategy allows imaging single-molecule dynamics of proteins involved in intraflagellar transport (IFT), a transport mechanism specific to cilia (e.g. kinesin-2, cytoplasmic dynein-2, etc. [19, 21, 23, 24]) as well as IFT-cargo proteins such as the Ca^2+^-channel OCR-2, which plays a key role in signalling [22]. We also obtained first insights in how the dendritic transport machinery transports vesicles containing OCR-2 from cell body, along dendrite, towards the base of the cilium. Although our approach has proven successful and insightful for several labeled proteins, we noticed that for some proteins the time during which single-molecule imaging in cilia or dendrite is possible is only brief, limited by the photobleaching of all molecules, severely reducing the amount of data that can be taken from a single sample.

**Figure 1:**
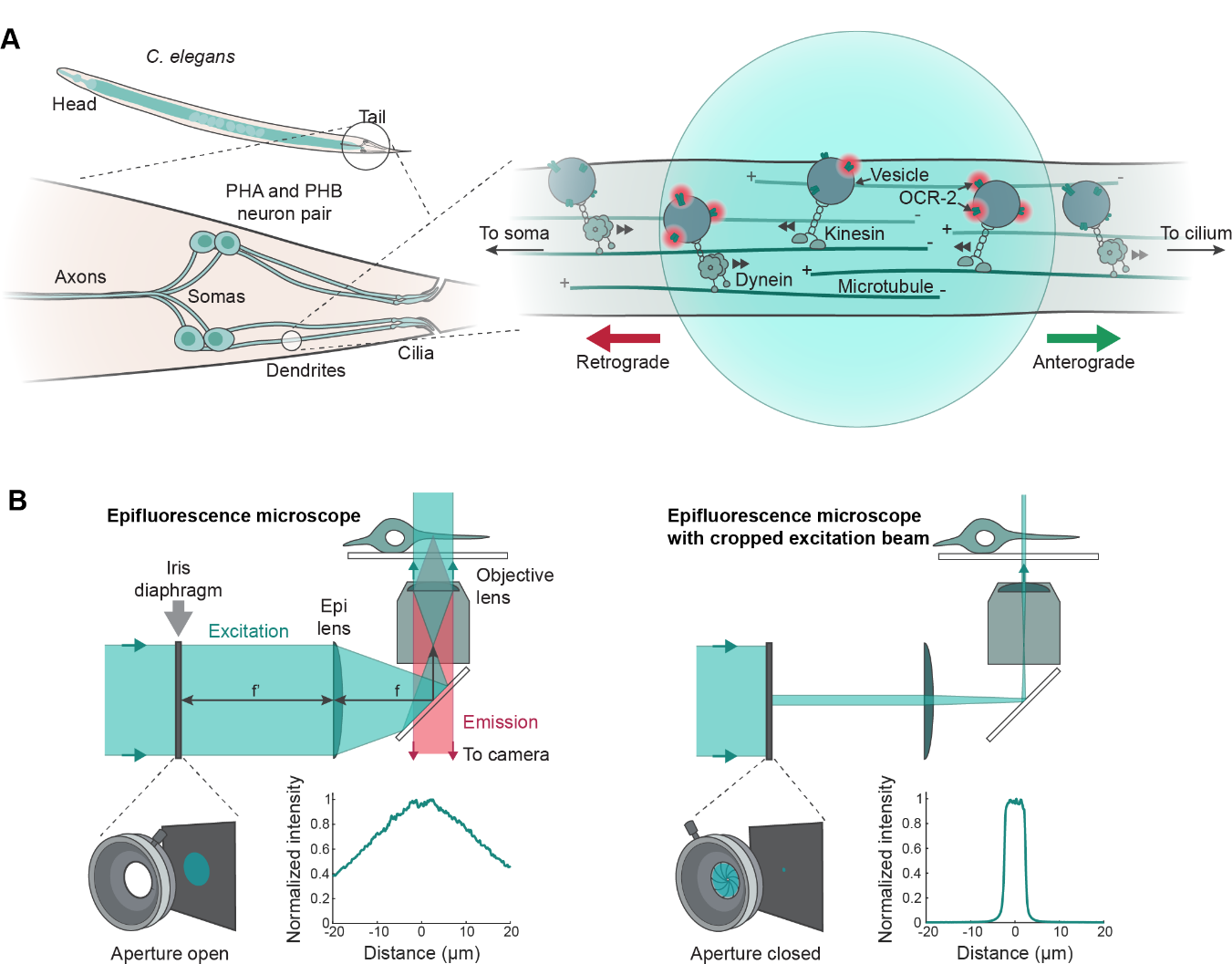
Illustration of small-window illumination microscopy (SWIM) and the imaged biological system. **(A)** Cartoon illustration of *C. elegans*, highlighting the ciliated chemosensory neurons, PHA and PHB, located in the tail of *C. elegans*. These neurons have dendrites which are tens of µm long, connecting the soma (cell body) to the cilia. Vesicles containing the transmembrane Ca^2+^ channel OCR-2 are transported in the anterograde (from the soma to the cilium) and retrograde (from the cilium to the soma) direction along microtubule bundles in the dendrites, driven by microtubule-based motors, dynein (anterograde) and kinesin (retrograde). The blue circle (illustration to the right) represents the region in the dendrite illuminated using SWIM, with fluorescently-labelled OCR-2 molecules being excited and fluorescing (indicated by red circles) upon entering the excitation region. **(B)** Schematic illustration of SWIM. The sample is imaged using laser-illuminated widefield epi-fluorescence microscopy, where the excitation beam size can be changed using an iris diaphragm at a focal plane conjugate to the image plane of the objective (at the back-focal plane of the epi lens, f). When the aperture of the diaphragm is open (left), the field of view is illuminated with a large parallel beam (beam width ∼30 µm, 2σ; see inset left plot). Closing the aperture (right) results in a cropped small-diameter excitation beam (window width ∼5-10 µm; see inset right plot).

We reasoned that we could prolong the time window in which single-molecule imaging is possible by only exciting a smaller region of the sensory neuron to allow continuous replenishment of ’fresh’, unbleached proteins into the imaged region. Here, we show that restricting the excitation area in the *xy*-plane (Figure 1B) – an approach we call Small-Window Illumination Microscopy (SWIM) – allows for considerably longer-duration single-molecule imaging *in vivo* in systems where proteins continuously move in and out of the excited area. Furthermore, remarkably, we discover that performing single-molecule imaging, using SWIM, results in a substantially higher SBR. Using careful experiments, numerical simulations and analytical calculations, we demonstrate that having a small excitation window minimizes autofluorescence and out-of-focus fluorescence, thereby improving the imaging quality. Finally, we utilize SWIM to reveal the dendritic transport dynamics of OCR-2-associated vesicles moving between the neuron cell-body (soma) and the cilium (Figure 1A).

## Results and Discussion

The typical diameter of the excitation beam in our set-up, as we have used in previous studies [19–24], is ∼30 µm (2*σ*, Gaussian; Figure 1B left), roughly covering the field of view of the EMCCD camera (41 x 41 um). To implement SWIM, we placed an iris diaphragm in the excitation path, which allows us to change the size of the excitation beam (Figure 1B). Closing the aperture of the diaphragm to the minimal of 0.8 mm yields an excited area with a diameter of ∼5 μm. The power density at the centre of the beam remains in the same range when cropping the beam with the diaphragm. To estimate the intensity profile of the excitation beam, with the aperture open or closed, we imaged a relatively uniform layer of Alexa555 dye, absorbed on a glass coverslip. When the aperture width is significantly smaller than the excitation beam, such that only the central part of the Gaussian beam passes through, the cropped excitation beam has an approximately rectangular profile, slightly smoothened near the edge (Figure 1B right and Supplementary Figure 1A). We observe that the intensity at the central part of the excitation beam is slightly lower when the aperture is closed (1.3x; Supplementary Figure 1A), suggesting a small optical loss due to cropping. Upon imaging reference fluorescent beads immobilized on glass, with the aperture open or closed (excitation window width ∼5 μm), we also observe a small drop in the signal intensity for beads near the centre of the illuminated window (0.95-1x; see Bead 1 and 2 in Supplementary Figure 1B-C), with the localization error increasing marginally (1-1.05x; Supplementary Figure 1D-F). Thus, placing an iris diaphragm in the excitation path allows to readily crop the excitation beam in order to excite a small region of interest in a sample, providing an evenly illuminated spot with a minor drop in beam intensity.

We utilized SWIM to image the transport dynamics of vesicles carrying the ciliary channel protein OCR-2 labelled with eGFP (OCR-2::eGFP), in the dendrites of the PHA/PHB neurons in *C. elegans* (Figure 1A). OCR-2, which is produced in the cell bodies, primarily localizes in cilia, which implies that it has to be transported across the ∼50 µm long dendrite that connects the soma to the cilium. Using a cropped 491 nm excitation beam, we imaged OCR-2::eGFP in a small region of a dendrite (excitation window width 7 µm) and observed rich dynamics of OCR-2-associated vesicles (Figure 2A and Supplementary Movie 1). Directed transport of vesicles both in the anterograde (soma to cilium) and retrograde (cilium to soma) direction, as well as purely diffusive vesicles, were observed. Although fluorescent proteins bleach in the illuminated region, due to the relatively high laser intensity, it is possible to image continuously for long durations (15 minutes of continuous imaging in Figure 2A), since new, unbleached proteins continuously move into the illuminated region. In contrast, illuminating with the full beam (aperture open) results in bleaching of fluorescing molecules over a much larger region, allowing observation of single OCR-2::eGFP for only a short period of time. Indeed, in the imaged dendrite shown in Figure 2A (Supplementary Movie 1), after opening the aperture following 15 mins of SWIM, to illuminate a much larger section of the worm (beam diameter ∼30 µm), OCR-2-associated vesicles could only be observed sparsely for 1-2 mins. In conclusion, SWIM allows imaging of the transport dynamics of OCR-2-associated vesicles in dendrites of *C. elegans* sensory neurons for considerably longer than is possible using standard widefield illumination.

**Figure 2:**
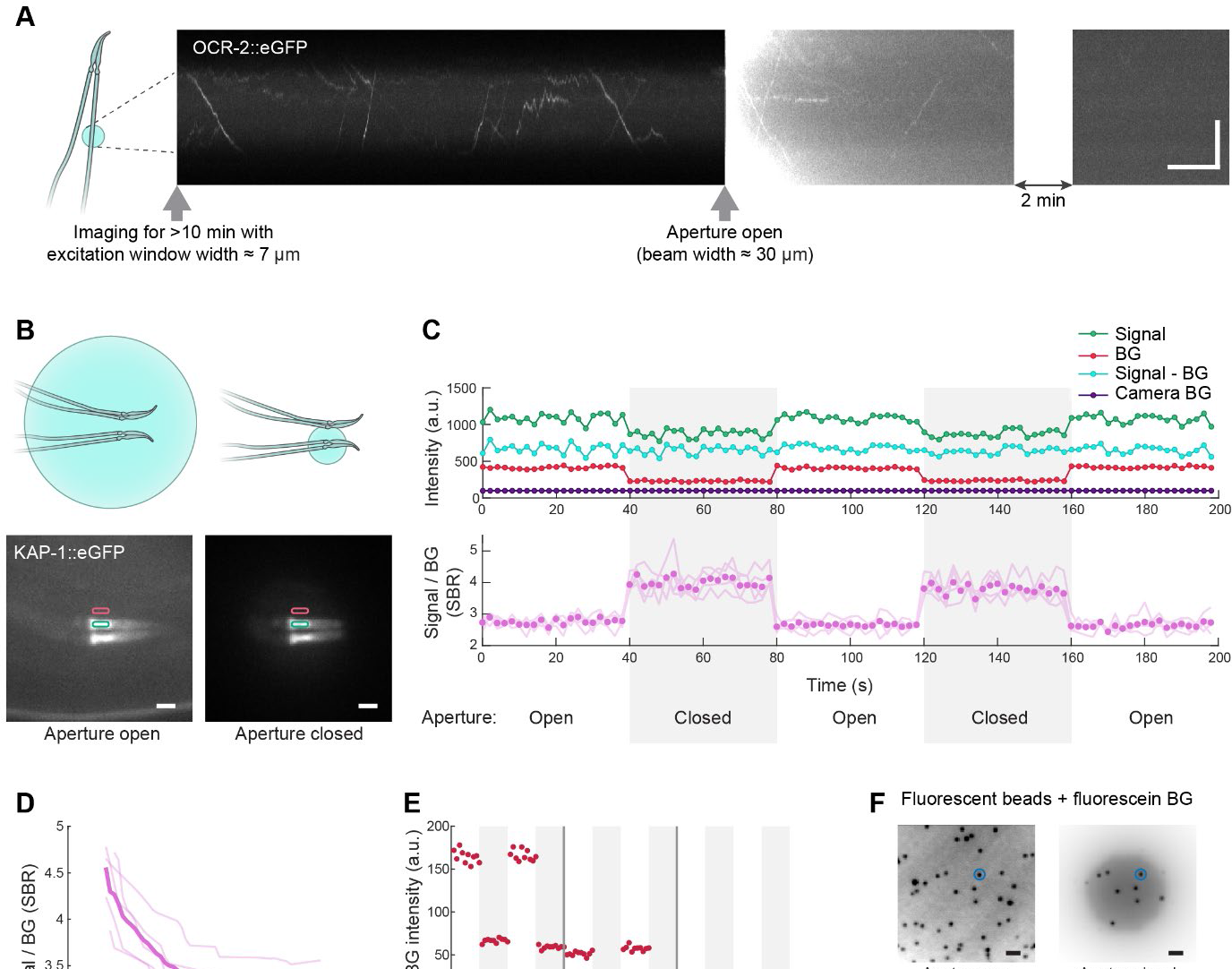
SWIM enables long-duration imaging and improved SBR (signal-to-background ratio) **(A)** Segments of a kymograph (time-intensity plot; timing of segments indicated) along a dendrite (also see **Supplementary Movie 1**), where OCR-2::eGFP-associated vesicles were imaged using a cropped 491 nm laser beam (excitation window width ∼7 µm; 6.67 fps). OCR-2-associated vesicles entering the small excitation window were continuously imaged for ∼15 mins. Upon opening the diaphragm at 15 mins, exposure by the entire beam (width ∼30 µm), quickly photobleached all OCR-2-associated vesicles within the larger excitation area and no more events were observed. Scale bars: 10 s (horizontal), 5 μm (vertical). Brightness and contrast are the same for all three segments. **(B)** Top: Schematic of imaging location (ciliary base) and approximate excitation window widths when the diaphragm aperture is open (left) and closed (right). Bottom: Time-averaged images of KAP-1::eGFP illuminated with aperture open (left) and closed (right). Regions, where the signal (green) and the BG (red) intensity were measured, are shown. Scale bar: 1 μm. **(C)** Top: Signal intensity (Signal), BG, their difference (Signal - BG), and the camera offset (Camera BG), measured over time in the regions indicated in B. Bottom: SBR as a function of time obtained from 5 worms (light magenta lines; mean indicated with circles). The aperture state is indicated below the x-axis. **(D)** Mean SBR obtained from 5 worms imaged at different excitation window widths (individual events plotted in light magenta; see details of example worm in Supplementary Figure 2C). The most drastic change in SBR happens at small window widths up to ∼10 μm. Dotted line indicates SBR (mean from 5 worms) when the aperture is completely open. **(E)** BG intensity imaged in a wild-type worm with aperture alternatingly opened (white region) and closed (grey region) every 20 s. Arrowheads indicate the points at which the sample is photobleached (illuminating with 491 nm laser at maximal power for 2 min, first time with aperture closed (excitation window size ∼4 µm), second time with aperture opened (beam width ∼30 µm). **(F)** Top: Inverted image of fluorescent beads, with fluorescein in solution acting as fluorescence BG, imaged with aperture open (left) and closed (right). Brightness and contrast are differently scaled for the two images. Bottom: Localizations of a bead circled in the top images, tracked over 1000 frames with aperture open (left) and closed (right), centred around its mean xy position. Localization error (standard deviation, SD) is >2x higher with aperture open (also see Supplementary Figure 2D-E).

To our great surprise, the BG intensity appeared to be significantly lower when a smaller area was excited using SWIM (Figure 2A and Supplementary Movie 1). We sought to quantitatively characterize the effect of the excitation window width on the BG intensity. Since vesicles contain a varying amount of OCR-2, their intensities vary considerably, which makes quantitative comparison of SBR difficult. To overcome this, we imaged the relatively constant pool of KAP-1::eGFP (the non-motor subunit of kinesin-II) in the cilia of PHA/PHB neurons (Figure 2B, Supplementary Figure 2A) [21, 23]. First, we imaged the ensemble fluorescence intensity of KAP-1 in the cilia, performing time-lapse imaging (acquisition every 2 s) with low-intensity 491 nm excitation, switching the aperture between open and closed states every 40 s (Figure 2B). We measured the signal and BG fluorescence intensity and observed that while both signal and BG intensity decrease upon closing the aperture, the BG-corrected signal intensity is relatively independent of the aperture state (Figure 2C upper panel). This results in a substantial increase of the SBR from 3 to 4 upon closing the aperture (Figure 2C lower panel). We performed a similar experiment on single-molecules of KAP-1, using a high-intensity 491 nm excitation laser beam (Supplementary Figure 2A), switching the aperture between open and closed state every 15 s. Again, we observed that signal and BG intensities decrease upon closing the aperture, with SBR increasing by a factor of 1.5 (Supplementary Figure 2B). Thus, exciting only a small region in a sample considerably improves the quality of single-molecule imaging by increasing the SBR of individual molecules. Next, we explored how the SBR depends on the width of the excited area, by imaging the bulk fluorescence intensity of KAP-1 at different aperture widths (Supplementary Figure 2C, Figure 2D). We discovered that the SBR decreases non-linearly with increasing window size, changing rather sharply for small window widths (5–15 µm). In principle, it should be possible to increase the SBR further by using even smaller excitation windows. Exciting larger areas (>20 µm) results in the SBR saturating to the value obtained with the full beam. Taken together, it appears that while the BG-corrected signal intensity remains constant, the BG intensity decreases with the reduction in size of the excited region, resulting in the significantly higher SBR.

Since the BG intensity crucially determines the quality of single-molecule imaging we wanted to better understand its source. To this end, we imaged in the tail region of a worm not expressing any fluorescent proteins (and thus only producing BG light upon excitation) using low-intensity 491 nm laser excitation, while periodically opening and closing the aperture. We observed an approximately 70% reduction of the measured BG light intensity in the central region of the sample with the aperture closed (Figure 2E, left). We repeated these measurements (on the same area of the same worm) after prolonged high-intensity 491 nm illumination with the aperture closed, thereby photobleaching only the region illuminated by the small excitation window (Figure 2E, middle), and observed that the light intensity nearly dropped to zero with the aperture closed, while it was still substantial with the aperture open. Subsequently, we illuminated the sample with high-intensity 491 nm laser light with the aperture open, photobleaching the entire region, and repeated the measurement cycle (Figure 2E, right). We now detected only a very small light intensity, both with the aperture closed and open. From these measurements we can draw two conclusions regarding BG light signals produced when exciting live *C. elegans* with 491 nm laser light. (I) The BG signal is mostly due to autofluorescence in the body of the worm, since it nearly disappears after prolonged, high-intensity excitation. The small residual BG after photobleaching might be due to elastic or inelastic (Raman) scattering of the excitation light [25, 26], or incomplete photobleaching. (II) When illuminating with the aperture open, a significant fraction of the BG signal at the central region of the focal plane comes from the part not illuminated with the cropped excitation beam. This is evident from the large BG observed upon excitation with the aperture open after photobleaching with the aperture closed (Figure 2E middle). Finally, we imaged fluorescent beads immobilized on glass with fluorescein in solution, to provide BG fluorescence, with aperture open and closed (Figure 2F, Supplementary Figure 2D-E). Localization error of the beads improves by a factor of 2x with aperture closed. These experiments clearly indicate that the intensity of the out-of-focus autofluorescence decreases substantially when limiting the excitation area in epifluorescence microscopy.

To provide a theoretical basis to this observation that the autofluorescence BG decreases when using a smaller excitation window, we performed (analytical) calculations, with the following assumptions. First, we assume that our excitation beam is Gaussian, characterized by a beam diameter *B*_*d*_ (= 2*σ*_*exc*_; with *σ*_*exc*_ the standard deviation). Furthermore, we assume that autofluorescence is generated homogeneously in the sample and that its local intensity is proportional to the excitation intensity. We assume out-of-focus light to propagate through our sample as a high-NA focused Gaussian beam (illustrated in Figure 3A). This means that out-of-focus autofluorescence generated in a point at height *z* will generate a Gaussian-shaped beam in the focal plane (*z*=0). The width of this beam ((*z*)) depends on *z*, the wavelength of the fluorescence, *λ*, and sin *θ* = *NA*/*n* (with *θ* the half acceptance angle of the objective, NA is the numerical aperture and *n* is the refractive index of the immersion medium), following [27]:

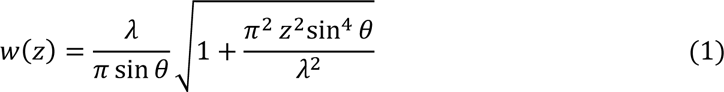

**Figure 3:**
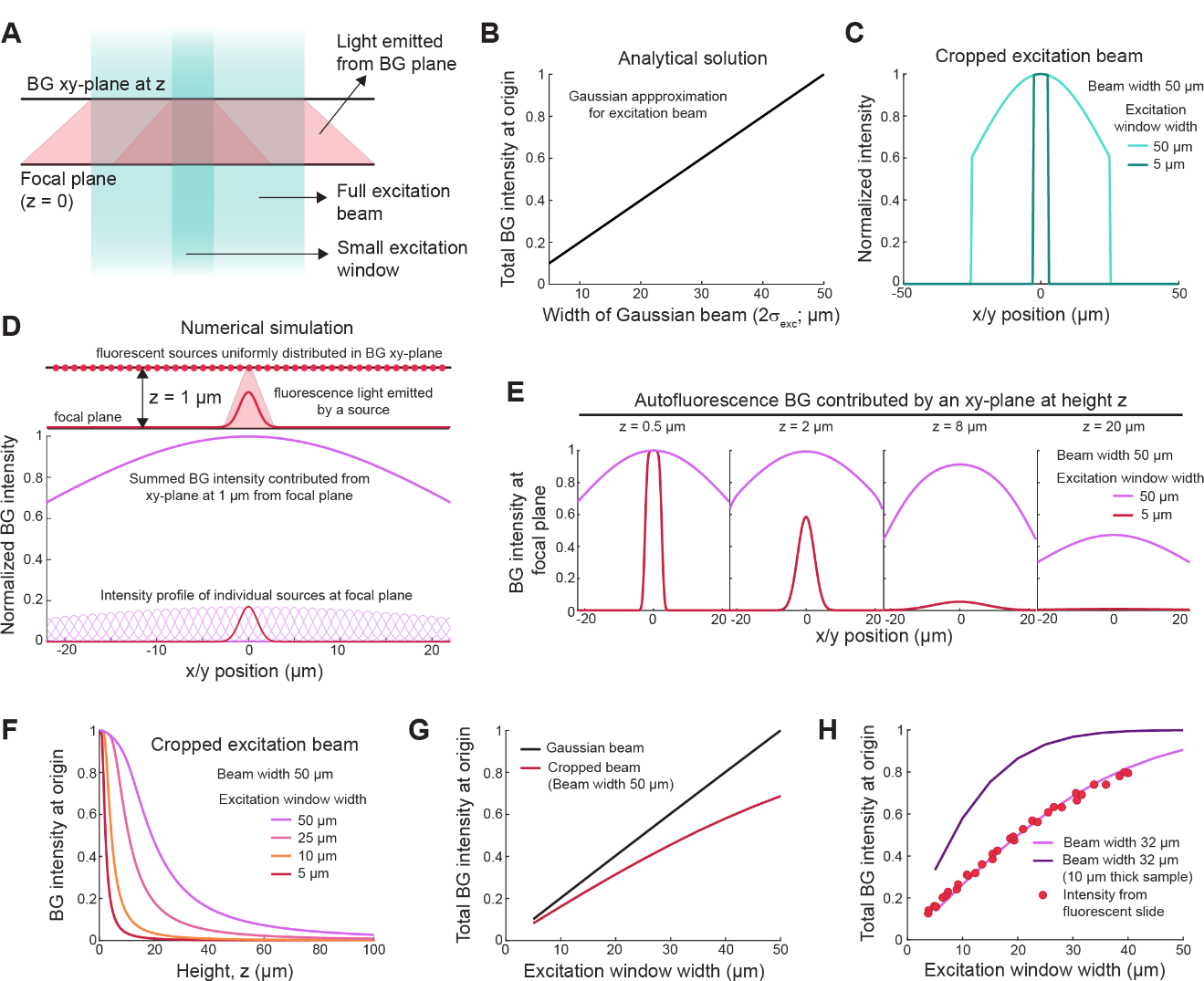
Computer simulations explain the change in BG autofluorescence with the width of the excitation beam. **(A)** Schematic representation of excitation beams with different widths, (cyan shaded area) propagating through a sample. BG autofluorescence light is emitted from an out-of-focus BG xy-plane at height z and propagates through the sample (red shaded area). **(B)** Analytical calculation assuming parallel Gaussian excitation beam shows that the BG intensity from the whole sample detected at the origin (x = y = z = 0 μm) increases linearly with the width of the excitation beam. **(C)** The normalized intensity profile of a Gaussian excitation beam (beam width 50 μm), cropped by a diaphragm with aperture width 50 μm (light cyan) and 5 μm (dark cyan), ignoring diffraction effects. **(D)** Illustration of numerical simulations that shows an example BG xy-plane at z = 1 μm, that contains a uniform and continuous distribution of fluorescent sources. When excited, each source emits light, propagating through the sample as a high-NA Gaussian beam with focus at the location of the source. The emission intensity of each source is proportional to the excitation intensity at the location of the source. The total BG intensity passing through the focal plane, originating from sources at BG xy-plane at height z, is obtained by summing the intensities of all individual sources in that plane. **(E)** BG intensity contributed by individual xy-planes (at 0.5 μm, 2 μm, 8 μm and 20 μm) illuminated by a Gaussian excitation beam (beam width 50 μm) cropped by apertures with width 50 μm (magenta) or 5 μm (red). **(F)** Relative contribution to BG at the origin, from xy-planes at different height (z) with respect to the focal plane (z=0). Sample illuminated by an excitation beam (width 50 μm) cropped by apertures of indicated widths. **(G)** Total BG intensity at the origin as a function of beam width (for illumination with Gaussian excitation beam of increasing beam widths; black line) or aperture width (for illumination with a 50 μm Gaussian excitation beam cropped by an aperture of increasing widths; red line). **(H)** Total BG intensity at the origin as a function of aperture width (illumination with a cropped excitation beam with full width 30 μm), scanning through a thick sample (0 to 1 mm; magenta line) and a thin sample (0 to 10 μm; dark purple line). The red datapoints represent the experimentally determined intensity at the centre of a fluorescent plastic slide (of thickness ∼1.6 mm) for different aperture widths, using a cropped excitation beam with a full width of ∼30 μm. In the whole figure the 2D intensity profiles used in our analytical calculations and numerical simulations are represented by 1D cross-sections, for ease of visual representation.

The autofluorescence propagating from excitations in an *xy*-plane at height *z*, generated by the Gaussian excitation beam in the focal plane, will then be the convolution of the Gaussian excitation beam with the high-NA focused Gaussian emission beam, creating a Gaussian profile with normalized intensity:

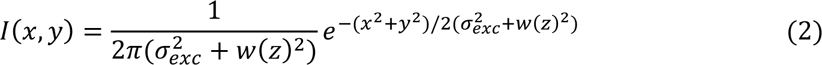

To calculate the BG intensity contribution of different out-of-focus planes for different widths of the excitation beam (*I*_*bg*_(*z*, *σ*_*exc*_)), we consider the intensity at the origin (*x*=*y*=*z*=0), where the exponential term is equal to one and only the pre-factor counts. We normalize the intensity by dividing the contribution from an *xy*-plane at height *z* by the contribution from the focal plane (*z*=0), obtaining the following:

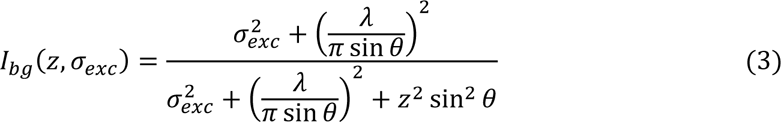

This equation can be readily integrated over *z* (from minus to plus infinity):

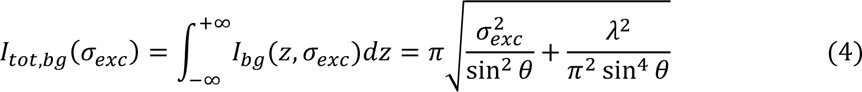

This can be further simplified realizing that our model for the parallel Gaussian excitation beam is only valid for beam diameters much larger than the diffraction limit. In that case, the second term in the square root is negligible, resulting in:

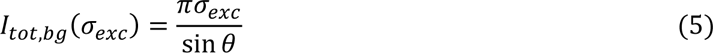

In this equation, the autofluorescence BG intensity increases linearly with the width of the excitation beam, assuming the excitation beam is a parallel Gaussian (Figure 3B). In our experiments, however, we use an aperture to crop an (approximately) Gaussian beam with a constant beam width, which results in a beam profile with a different shape than considered in the analytical calculation (see Figure 3C). To explore how the autofluorescence BG intensity depends on the width of a Gaussian excitation beam (of constant beam width 50 μm) that is cropped by varying the aperture size, we performed numerical simulations. Again, we assume that the sources of autofluorescence in our sample are uniformly and continuously distributed. Autofluorescence generated at a given point in the sample propagates from this point as a high-NA Gaussian beam, resulting in a beam width at the focal plane given by equation (1) and has an amplitude proportional to the intensity of the cropped excitation beam at that point. Summing up the autofluorescence BG intensity contributions at *z* = 0, from all points in an *xy*-plane at height *z*, gives the BG intensity at the focal plane, emanating from the plane at *z* (Figure 3D). We found that the autofluorescence BG intensity at the origin (*x*=*y*=*z*=0) contributed by an out-of-focus plane at *z* decreases with increasing *z* and does this much more steeply for a more cropped excitation beam (Figure 3E and 3F). The total autofluorescence BG at the origin increases with increasing aperture width, saturating when the aperture width becomes similar in size to the width of the full Gaussian beam (red line in Figure 3G). We also performed similar numerical simulations assuming that the excitation beam is a Gaussian with varying width (Supplementary Figure 3A-C), and found that the numerical simulation gives the same result as the analytical calculation (using the same assumptions; black line in Figure 3B and 3G). To experimentally test these theoretical predictions, we measured the fluorescence intensity at the centre of a fluorescent slide (of thickness ∼1.6 mm) excited by the excitation beam used in our experiments, cropped by an aperture with different widths. We measured an increase in BG autofluorescence with increasing aperture widths (Supplementary Figure 3D), quantitatively very similar to the numerical simulations (Figure 3H), indicating that our theoretical model explains that SWIM results in a substantially improved SBR because of reduction of the out-of-focus autofluorescence background. When the sample is relatively thin (for instance 10 μm), the SBR can only be improved by employing a substantially cropped excitation beam (< 20 μm; blue line in Figure 3H). This is likely the case in our *C. elegans* experiment. At the location where we image (in the tail), the worm is ∼10-20 μm thick. It is thus not surprising that we observe that the SBR does not decrease further for aperture widths > 20 μm (Figure 2D).

In summary, the key to performing single-molecule imaging in neurons of living *C. elegans* is to limit the number of fluorescent molecules of interest and to minimize BG autofluorescence. Both can be achieved with the relatively high excitation laser intensities used for single-molecule imaging. We have shown here that the BG (primarily due to out-of-focus autofluorescence) can be substantially decreased when only a small region is excited using SWIM. In addition, SWIM allows for longer duration single-molecule imaging, because fresh, unbleached fluorescent molecules of interest can continuously move into the illuminated region of interest. Longer duration imaging not only improves data throughput, it also helps to further reduce the BG thanks to photobleaching by the excitation light (as seen in Supplementary Figure 2A-B).

Having established that reducing the excitation area can substantially improve single-particle imaging *in vivo*, we utilized SWIM to image the dynamics of OCR-2-associated vesicles in the dendrites of phasmid neurons in the tail of *C. elegans* (illustrated in Figure 4A). As seen in the example movie (Supplementary Movie 2; image acquisition rate 6.67 fps), it was possible to visualize OCR-2::eGFP in a section of both the PHA and PHB dendrites (denoted dendrite 1 and 2 in Figure 4B and Supplementary Movie 2, since we cannot assign them to either PHA or PHB), since both dendrites were in one Z-plane. We observed that OCR-2-associated vesicles are continuously being transported, in a directed manner, across the imaged region of the tens of µm long dendrites, from the soma to the ciliary base (anterograde) and back (retrograde). OCR-2, which is a subunit of a TRPV channel associated with sensory cilia [28], is expressed in the neuron’s soma and associates with vesicles formed by the trans-Golgi network [22], that likely constitute the anterograde moving vesicles. The retrograde-directed movement of OCR-2 from the cilia towards the soma was slightly less anticipated. Since OCR-2 primarily localizes at the membrane of the cilia and the periciliary membrane compartment (PCMC), at the transition between dendrite and cilium, we hypothesize that retrograde-moving OCR-2 molecules are associated with endocytic vesicles formed at the PCMC [29], cycling old OCR-2 proteins back to the soma. Further, we observed that a significant portion of the vesicles was purely diffusive, typically photobleaching within the first few seconds of imaging (Figure 2A and Supplementary Figure 5A).

**Figure 4:**
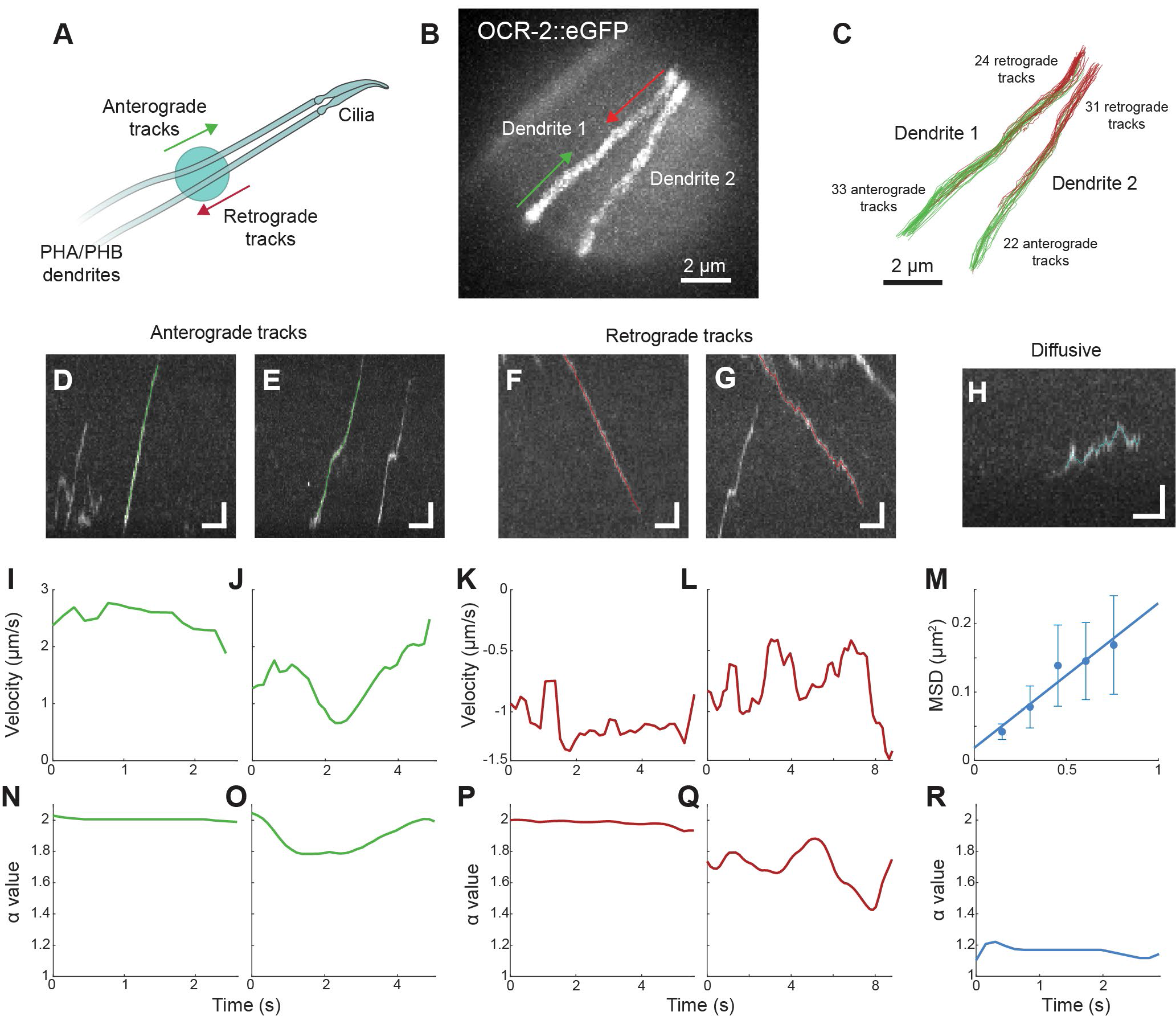
Single-particle tracking of OCR-2-associated vesicles in dendrites, imaged using SWIM. **(A)** Illustration of SWIM in dendrites of phasmid neurons (left). **(B)** Maximum projection of an example movie (also see **Supplementary Movie 2;** imaged using 491 nm laser at 6.67 fps) shows the section of PHA and PHB dendrites (labelled dendrite 1 and dendrite 2) illuminated by a small excitation window (window width ∼11 µm), where the dynamics of OCR-2:eGFP associated with vesicles can be visualized. **(C)** Single-particle tracking in the example movie yields 33 anterograde (soma to cilium) and 24 retrograde (cilium to soma) tracks in dendrite 1, and 22 anterograde and 31 retrograde tracks in dendrite 2. **(D-H)** Kymographs (from dendrite 1) overlayed with track data showing typically observed anterograde (D-E; green tracks), retrograde (F-G; red tracks) and diffusive (H; blue track) events. Horizontal scale bar: 2 s; Vertical scale bar: 1 μm. **(I-L)** Point-to-point velocity plotted over time, for the anterograde (I-J) and retrograde (K-L) tracks calculated for the events shown in D-E and F-G. **(M)** Diffusion coefficient for the diffusive event shown in H calculated using MSD analysis is 0.11 μm^2^/s. **(N-R)** Alpha value, calculated using a windowed mean-square displacements-based approach, plotted over time, for the anterograde (N-O) retrograde (P-Q) and diffusive (R) tracks corresponding to the events shown in D-H. Alpha value is ∼2 for purely directed transport and ∼1 for purely diffusive transport.

We performed single-particle tracking to obtain more detailed information on the transport dynamics (Figure 4C). Both retrograde and anterograde trajectories appeared, in some cases, entirely directed (e.g. Figure 4D and 4F) and in others, directed stretches where interspersed with diffusive phases (e.g. Figure 4E and 4G). Diffusive vesicles (e.g. Figure 4H) were mostly observed only briefly, mostly near the edges of the excitation area, getting photobleached quickly or moving out of the excited area. We analysed the local velocities of individual trajectories (referred to as point-to-point velocity) by calculating the local slope from the distance-time data (including 3 consecutive frames). For entirely directed anterograde tracks, the velocity is ∼2-2.5 μm/s (Figure 4I), while for less purely directed trajectories, the velocity is lower in the more diffusive sections (∼1-2 μm/s; Figure 4J). Similarly, for entirely directed retrograde trajectories the velocity is ∼1-1.2 μm/s (Figure 4K; negative value indicates retrograde motion), while for less purely directed tracks, velocities range between ∼0.5-1 μm/s (Figure 4L). For diffusive trajectories, we calculated the diffusion coefficient from the mean squared displacement (MSD) to be 0.11 μm^2^/s (for the trajectory of Figure 4H; Figure 4M). To obtain a quantitative measure for the directedness of the motion, we used an MSD-based approach [22, 23, 30] to extract the anomalous exponent (α) from *MSD*(*τ*) = 2Γ*τ*^*α*^ (where Γ is the generalized transport coefficient and *τ* is the time lag) along the track, in the direction of motion. α is a measure of the directedness of the motion, α = 2 for purely directed motion, α = 1 for purely diffusive motion and α < 1 for subdiffusion or pausing [31]. We found that α for apparently entirely directed trajectories (both anterograde and retrograde) is 2 (Figure 4N and 4P), while it ranges between 1.5 and 2 for less directed tracks (Figure 4O and 4Q), and is close to 1 for diffusive trajectories (Figure 4R).

Since directed trajectories displayed some variability in motion, we filtered directed regions of individual trajectories based on the α values, to obtain a better estimate of the velocity of the molecular motors driving the motion. For anterograde trajectories, the average point-to-point velocity from 144 tracks (N = 2865) in 7 dendrites (5 worms) was 1.77 ± 0.05 μm/s (Figure 5A; average value and error estimated using bootstrapping; see methods). Using a threshold of α = 1.95, we find that less directed stretches of trajectories (α < 1.95) show an average velocity of 1.58 ± 0.07 μm/s (N = 1694) and highly directed stretches of trajectories (α > 1.95) show an average velocity of 1.99 ± 0.06 μm/s (N = 1171; Figure 5A). Binning the time points in 0.05 α-value bins, shows that the velocity increases from 1 μm/s (α: 1.5-1.55) to 2.1 μm/s (α: 2-2.05; Supplementary Figure 4A). For retrograde tracks, the average point-to-point velocity from 184 tracks (N = 6375) in 8 dendrites (5 worms) is 0.82 ± 0.01 μm/s (Figure 5B). Furthermore, we find that less directed stretches of trajectories (α < 1.95) show an average velocity of 0.75 ± 0.02 μm/s (N = 4662) and highly directed stretches (α > 1.95) show an average velocity of 0.95 ± 0.02 μm/s (N = 1805; Figure 5B). Binning the velocity data using α values, we find that the velocity increases from 0.55 μm/s (α: 1.5-1.55) to 0.95 μm/s (α: 2-2.05; Supplementary Figure 4B). 40% of the tracked datapoints for anterograde tracks and 28% of the tracked datapoints for retrograde tracks have α > 1.95, indicating that the OCR-2-associated vesicles in the anterograde direction are less diffusive. Since the microtubule bundles in the dendrites of *C. elegans* neurons are minus-end out [32–34], anterograde vesicles are most likely driven by cytoplasmic dynein-1 [35] and retrograde vesicles are likely driven by kinesin-1 (UNC-116) and/or kinesin-3 (UNC-104) [36, 37]. Thus, using filtering based on α values, we find that in dendrites of sensory neurons in *C. elegans*, cytoplasmic dynein drives OCR-2 vesicles at 2.1 μm/s (slightly higher than previously observed [38], and kinesin drives retrograde OCR-2 vesicles at 0.95 μm/s (similar to values reported for kinesin-3 [36], but not kinesin-1 [38]).

**Figure 5:**
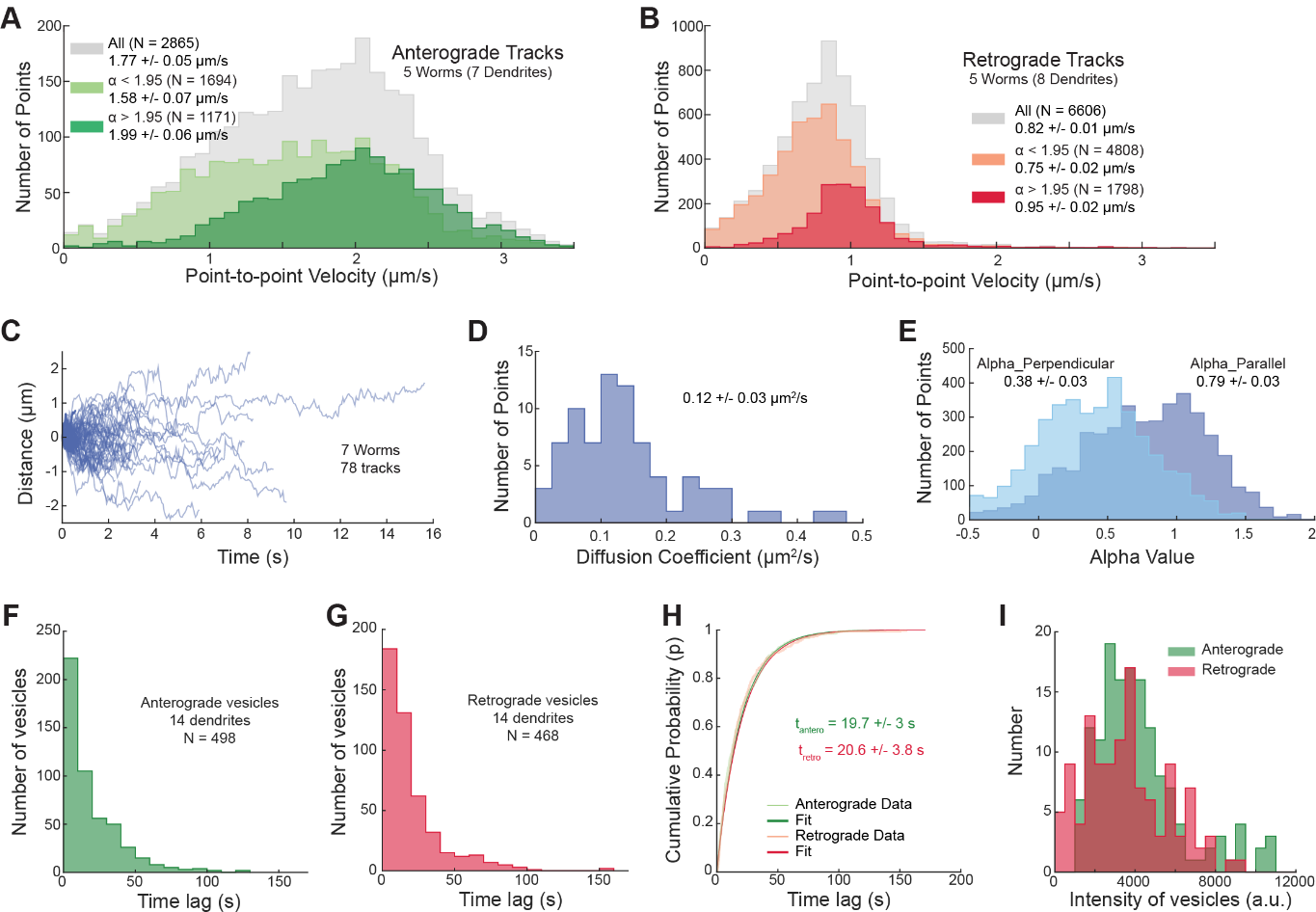
Statistical analysis of the motion of OCR-2-associated vesicles moving in and out of the illuminated region. **(A)** Histograms of point-to-point velocities obtained from 277 tracks of OCR-2-associated vesicles moving in the anterograde direction, in 7 dendrites (5 worms). Average velocity obtained from all data points is 1.77 ± 0.05 μm/s (N = 2865; grey). For data points with α < 1.95, average velocity is 1.58 ± 0.07 μm/s (N = 1694; light green), while, for data points with α > 1.95, it is 1.99 ± 0.06 μm/s (N = 1171; green). **(B)** Histograms of point-to-point velocities obtained from 228 retrograde tracks, in 8 dendrites (5 worms). Average velocity obtained from all data points is 0.82 ± 0.01 μm/s (N = 6606; grey). For data points with α < 1.95, the average velocity is 0.75 ± 0.02 μm/s (N = 4808; light red), while, for data points with α > 1.95, it is 0.95 ± 0.02 μm/s (N = 1798; red). **(C)** Distance-time plots of 78 tracks of diffusive OCR-2-associated vesicles, in 7 dendrites (7 worms). **(D)** Histogram of diffusion coefficients of the tracks in C, with average diffusion coefficient 0.12 ± 0.03 μm^2^/s. **(E)** Histograms of the α-values in the direction perpendicular (light blue; 0.38 ± 0.03; N = 4028) and parallel (dark blue; 0.79 ± 0.03) to the length of the dendrite. Average value and error are estimated using bootstrapping. **(F)** Histogram of the time lag between subsequent OCR-2-associated vesicles moving in the anterograde direction, in 14 dendrites (N = 498; 9 worms). **(G)** Histogram of the time lag between subsequent OCR-2-associated vesicles moving in the retrograde direction, in the same 14 dendrites (N = 468). **(H)** The time-lag distribution in F and G plotted as a cumulative distribution function, overlayed with the least square fit to the function, in green and red for the anterograde and retrograde vesicles, respectively. The characteristic time between subsequent anterograde OCR-2 vesicles is 19.7 ± 3 s and between subsequent retrograde OCR-2 vesicles is 20.6 ± 3.8 s. **(I)** Histograms of intensities of anterograde (N = 146; 6 dendrites in 4 worms) and retrograde (N = 122; 8 dendrites in 5 worms) vesicles from two dendrites in the same worm. The average intensity of anterograde vesicles was 3771 ± 451 a.u. while the average intensity of retrograde vesicles was 3399 ± 670 a.u. Average value and error are estimated using bootstrapping.

As described above, some vesicles appear entirely directed, others switch between directed and less directed (having a diffusive component) phases, leading to a substantial variability in speeds. In rare cases, reversal of motion from anterograde to retrograde, or *vice versa*, was also observed (Supplementary Figure 5C), suggesting that OCR-2-associated vesicles can associate with retrograde and anterograde directed motors simultaneously. It is possible that directionality and directedness are tuned by the number and ratio of activated dynein and kinesin motors cooperating to drive a vesicle [39]. Trajectories of vesicles driven by a large number of motors with the same directionality would show highly directed motion. Less directed motion might be observed when a trajectory is due to a vesicle driven by only a few motors of the same directionality. Because of the finite run length of the motors, a vesicle might regularly switch to a transient diffusive state whenever none of the motors connects it to the microtubule lattice. Another cause of less directed trajectories might be vesicles containing opposite directionality motors (i.e., both dynein and kinesin), resulting in transient directed runs, interspersed by periods of tug-of-war or reversals of direction [39, 40].

Next, we explored the nature of the diffusive trajectories. To obtain diffusion coefficients of the diffusive vesicles we tracked 78 events in 7 dendrites (Figure 5C). The distribution of diffusion coefficients for individual tracks, is broad with an average diffusion coefficient of 0.12 ± 0.03 μm^2^/s (Figure 5D). Remarkably, the diffusivity of single OCR-2 molecules along the ciliary membrane is much lower (∼0.03 μm^2^/s [22]). For these trajectories, the average α along the length of the dendrite is 0.79 ± 0.03 and the average α perpendicular to the dendrite longitudinal axis is 0.38 ± 0.03 (Figure 5E), indicating that the motion of the vesicles is slightly sub-diffusive, perhaps due to transient interactions with the microtubule cytoskeleton. We also observed that anterograde and retrograde vesicles occasionally switched to a diffusive state (Supplementary Figure 5B), the mechanism of which is unknown but could be linked with the disassociation or deactivation of the driving motors. We did not observe diffusive vesicles switching to directed motion. This might reflect, however, a limitation of our imaging approach, in which the diffusive vesicles are mostly visualized near the edge of the excitation window, moving back and forth, before they eventually photobleach.

Finally, SWIM makes it possible to estimate the rate at which OCR-2 proteins move back and forth along an imaged dendritic section. We determined that the distribution of time intervals between individual anterograde (N = 498 in 14 dendrites; Figure 5F), as well as retrograde (N = 468 in the same 14 dendrites; Figure 5G) OCR-2-associated vesicles is exponentially distributed. The characteristic time interval between two anterograde vesicles was 19.8 ± 3.2 s and between two retrograde vesicles was 20.6 ± 4 s (Figure 5H), suggesting that anterograde and retrograde vesicles have similar frequencies. Furthermore, we quantified the intensity of the vesicles and observed a broad distribution of intensities, indicating that the amount of OCR-2 associated to individual vesicles is highly variable (Figure 5I). We also observed that the intensity of the retrograde vesicles (3399 ± 670 a.u.) was on average slightly lower than the intensity of anterograde vesicles (3771 ± 451 a.u.; Figure 5I). This may imply that slightly more OCR-2 proteins move towards the cilium than away from it (to the soma). It is plausible that only part of the OCR-2 returns back to the soma, while the rest is either degraded or extruded out from the cilia, as part of extracellular vesicles [41]. In summary, employing SWIM and single-particle analysis, we provide a detailed, quantitative picture of the dynamics of vesicles transporting OCR-2 proteins along the dendrites of *C. elegans* sensory neurons.

## Conclusion

In this work, we have developed a simple technique, referred to as small-window illumination microscopy (SWIM), to perform single-molecule fluorescence imaging for long durations (at least 15 minutes) in sensory neurons of living *C. elegans* worms. In SWIM we remarkably improve the quality of single-molecule imaging by restricting the size of the excitation area in the *xy*-plane in order to reduce the autofluorescence BG. In our implementation of SWIM, we used an iris diaphragm in the excitation path to reduce the width of the region excited in the sample, mostly because of ease of implementation in our existing setup and to create a more or less flat-top excitation profile. In an alternative approach, a purely Gaussian beam could be expanded (or compressed) using a variable beam expander, limiting diffraction or reflection artifacts arising from the iris, but at the cost of a large intensity variation around the centre of the beam. For removing out-of-focus fluorescence in the z-direction, techniques such as TIRF, HILO or light sheet microscopy are superior to SWIM, but for long duration imaging in a small spatial window, such approaches could be combined with SWIM. An alternative approach would be to scan a confocal excitation beam over the sample, fast, such that the whole region of interest would be scanned once (or more often) per camera frame. By employing a tightly focussed confocal excitation beam the BG autofluorescence could be further decreased, because of the decrease of the excitation intensity outside of the image plane. Thus, while our approach of cropping an excitation beam might be the simplest means of implementing SWIM, there are other approaches that could further improve the quality of single-molecule imaging using a camera as detector.

We demonstrated the potential of SWIM by providing a detailed view on the dynamics of OCR-2-associated vesicles in the dendrites of chemosensory neurons in *C. elegans*. The optimal size (and shape) of the excitation window might be different for other applications. In principle, a smaller excitation window results in lower BG autofluorescence, but other factors such as optical properties of the fluorophore, the speed and nature of the transport dynamics, the density of labelled molecules, and the size of the region needed to be imaged, also play a role in determining the optimal excitation window width. In our view, SWIM is well suited for longer-duration single-molecule imaging in *in vivo* systems such as axons, dendrites, and cilia where fluorescently labelled proteins continuously move in and out of a given area. The dynamics of proteins in such systems are often highly heterogeneous, involving directed and diffusive transport, pausing, directional reversals, velocity variability, switching between modes of transport and tug-of-war behaviour [32, 40, 42-44]. SWIM can be of great benefit to address open biophysical questions in such *in vivo* systems.

## Methods

### C. elegans strains

*C. elegans* strains used in this study are listed in Table S1. Strains expressing eGFP-labelled OCR-2 and KAP-1 were generated earlier in our lab using CRISPR-Cas9 and Mos-1-mediated single-copy insertion, respectively [45, 46]. Worms were grown according to standard procedures [47], at 20°C, on nematode growth medium (NGM) plates seeded with HB101 *E.coli* bacteria.

### Fluorescence microscopy

Images were acquired using a custom-built epi-illuminated widefield fluorescence microscope [10, 21]. In short, the setup was built around an inverted microscope body (Nikon Ti E) equipped with a 100x oil immersion objective (Nikon, CFI Apo TIRF 100x, N.A.: 1.49). The system was controlled using MicroManager software (v1.4) [48]. Excitation light was provided by 491 nm DPSS laser (Cobolt Calypso, 50 mW). Laser power was adjusted using an acousto-optic tuneable filter (AOTF, AA Optoelectronics). For modulating the beam size, an iris diaphragm (Thorlabs, SM1D12, ø 0.8-12 mm) was mounted between the rotating diffuser [21] and the epi lens, at a distance equal to the focal length of the latter. The aperture size of the diaphragm was adjusted manually. Fluorescence was separated from the excitation light using a dichroic mirror (ZT 405/488/561rpc; Chroma) and an emission filter (525/45; Brightline HC, Semrock). Images were recorded by an EMCCD camera (Andor, iXon 897). Excitation laser power was measured with both open and closed aperture states, placing a power meter sensor (Thorlabs) at the objective mount. Peak power density remained in the same range (±10%) for both cases (calculated assuming that the uncropped beam has a Gaussian profile with *σ* = 15 µm, and minimal excitation window diameter = 5 µm).

#### *In vivo* imaging

For imaging live *C. elegans*, young adult hermaphrodite worms were sedated in 5 mM levamisole in M9 buffer (22 mM KH_2_PO_4_, 42 mM Na_2_HPO_4_, 85.5 mM NaCl, 1 mM MgSO_4_), sandwiched between an agarose pad (2% agarose in M9 buffer) and a coverslip and mounted on a microscope, as described in [10]. The region of interest (either cilia or dendrites of phasmid neurons) was brought into focus and imaged with an acquisition rate of 6.67-50 fps. Single-particle imaging was performed using high intensity 491 nm excitation (∼10 W/mm^2^), for bulk imaging the excitation intensity was >200 times lower. The Perfect Focus System (PFS) of the microscope was used to keep the sample in focus during long-duration experiments.

#### Control experiments

To accurately estimate the excitation beam profile, we coated the surface of glass with a thin layer of Alexa555 dye. Firstly, we labelled spermine with Alexa555 dye by reacting equimolar amounts of spermine with the corresponding NHS-ester of the dye in 50 mM borate buffer (pH 8.15) for one hour. Then, a solution containing 2 µl of 1% BSA and 0.4 µl of 140 µM Alexa555-spermine was incubated for a few minutes on a clean coverslip. The solution was then washed off and the remaining layer was imaged at ∼2-2.5% of the maximal power (∼0.2 W/mm^2^).

For measuring localization error and fluorescence intensity, fluorescent microspheres (Tetraspeck, 0.1 um; Invitrogen) diluted in M9 buffer were placed between a coverslip and a microscope slide, and imaged at ∼0.5% of the maximal power. For the localization error measurements with additional BG fluorescence, a mixture of fluorescent beads and fluorescein diluted with M9 buffer was flushed into a ∼0.1 mm high chamber sandwiched between a coverslip and microscope slide, with parafilm used as spacer.

To explore the effect of excitation beam size on measured autofluorescence BG intensity, ∼1.6 mm thick fluorescent plastic slides (Chroma) were imaged at ∼0.4% of the maximal power.

### Data analysis

#### Intensity analysis

Initial image analysis was performed in Fiji [49]. Ensemble and single-molecule fluorescence intensities were measured by manually selecting a region containing fluorescent signal and a region next to it within the illuminated area, for the background correction. The size of the excitation window was determined by drawing a circle around the region with highest intensity and determining its diameter (illustrated in Supplementary Figure 3D). Intensity profiles were made by manually drawing a line (width = 5 pixels) through the centre of the illuminated area and plotting the intensity along this line. In case of Alexa555-coated surface, to reduce intensity fluctuations due to inhomogeneities in the sample, for each aperture state the intensity was averaged over 10 independent locations in the sample and over 6 lines drawn through the centre of the illuminated region (with 60° angle between the lines).

#### Localization precision estimation

Fluorescent beads immobilised on the coverslip were tracked using a MATLAB-based software, FIESTA (version 1.6.0) [50] and the tracks were drift-corrected. Localization error was calculated as displacement standard deviation, 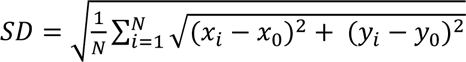, where *x*_0_ = *y*_0_ = 0 (mean *x*, *y* localization for each bead) and *N* is the number of tracked localizations.

#### Tracking of OCR-2 vesicles

OCR-2 vesicles were tracked using FIESTA. Tracks, corresponding to an event, contain information regarding time (*t_i_*), x and y coordinates (*x_i_*, *y_ii_*) and distance moved (*d_i_*), for every tracked time-frame *i*. The connected tracks were visualized and erroneous or small tracks (< 15 time-frames) were excluded from further analysis. Erroneous tracking primarily occurred when two or more retrograde, anterograde or diffusive events were too close to one another.

#### Point-to-point velocity for all tracks

Before calculating the point-to-point velocity, the tracks were smoothened by rolling frame averaging over 10 consecutive time frames, to reduce the contribution due to localization error (typically estimated to be between 10-80 nm, depending on the brightness of the tracked object). The point-to-point velocity was calculated using the following equation: *v_i_* = (*d_i+1_* − *d*_*i*−1_)/(*t*_*i*+1_ − *t*_*i*−1_).

#### Diffusion coefficient for diffusive tracks

Displacement along the length of the cilia for all diffusive tracks was calculated for discrete time points, and the displacement data were cumulated to calculate the average MSD (Mean Square Displacement) for every discrete time point. The first five points of the MSD thus obtained were then fitted with a linear curve, using the error bars as weights, with the slope of the fit providing the diffusion coefficient for each track.

#### Calculation of α-value

The x-y coordinates were transformed to dendritic spline coordinates, by defining a spline along the length of the imaged dendrite section and obtaining the distance parallel (*c*_||_*i*_) and perpendicular (*c*_⊥_*i*_) to the spline, for every tracked datapoint (*x*_*i*_, *y*_*i*_). The spline was generated by interpolating a cubic spline curve on a segmented line, drawn manually over all tracks in a given dendritic section (which provides the shape of the dendrite). For each datapoint (*c*_||_*i*_, *c*_⊥_*i*_), a windowed Mean Square Displacement classifier (wMSDc) approach, described in Danné et al. [30], was used to extract the anomalous exponent value (α) of the time lag (τ) from *MSD* = 2Γ*τ*^*α*^ (where Γ is the generalized transport coefficient), in the direction parallel (*α*_||_*i*_) and perpendicular (*α*_⊥_*i*_) to the spline. α was calculated analytically, using the following equation: *α* = 〈*α*〉 = 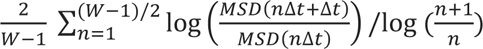; keeping a fixed window W = 15 time-frames. Due to the size of the window, all tracks shorter than 15 frames were removed from the analysis. In this paper, *α*_||_ values are referred to as α, since we primarily discuss the motion along the length of the dendrite and filter velocities based on *α*_||_ values.

#### Exponential fit for time-lag distribution

The time-lag between subsequent anterograde (or retrograde) vesicles was extracted by manually selecting events on a kymograph. The characteristic time lag (*τ*) was obtained by least-squares fitting of the cumulative distribution function (LSF-CDF), where the time lag distribution was used to generate a cumulative probability distribution that was fitted with the CDF: *y* = 1 − *e*^−*x*⁄*τ*^. The average value and error were estimated using bootstrapping (see below).

#### Vesicle Intensity Analysis

The intensity of a single vesicle was roughly estimated by manually drawing a box on the dendrite, with dimensions sufficient to capture all the light emitted by a single vesicle, positioned right at the edge of the illuminated spot, using Fiji. Because the square was placed at the edge of the illumination spot, we minimized prior excitation and photobleaching which would decrease the measured intensity. The intensity of actively transported vesicles passing through the square was recorded. The background intensity was measured by placing a box of the same dimension outside the dendrite and this was subsequently subtracted from the raw signal to obtain the background corrected value.

#### Estimating the average value and error for distributions

We utilized a bootstrapping approach to estimate parameters for any given distribution. The distribution (consisting of N measurements) was resampled by randomly picking N measurements from the measured distribution (with replacement) and calculating the median of the resampled distribution. This was repeated 1000 times. The resulting bootstrapping distribution was used to estimate the parameter (mean of the bootstrapping distribution *μ*) and its error (standard deviation of the bootstrapping distribution *σ*). All values and errors in this paper use *μ* ± 3*σ*.

#### Information on plots and figures

Kymographs were generated either on FIESTA or using the KymographClear [51] plug-in on Fiji/ImageJ. All the data was analysed and plotted using custom written scripts on MATLAB (The Math Works, Inc., R2021a).

### Numerical simulations to calculate the dependence of beam width on BG autofluorescence

To calculate the BG autofluorescence, we made the assumption that the excitation beam is parallel to and centred around the *z*-axis and has a Gaussian shape. For simulations where we change the size of the excitation window by varying the beam diameter, *B*_*d*_ (*B*_*d*_ = 2*σ*_*exc*_), of the excitation beam, the Gaussian intensity profile at a position (*l*_*x*_, *l*_*y*_) in an *xy*-plane is given by:

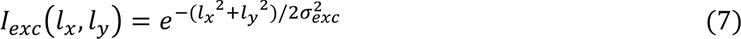

For the simulations where we change the size of the excitation window by cropping a Gaussian excitation beam of constant beam diameter *B*_*d*_ by changing the radius *r*_*a*_ of an aperture, the Gaussian intensity profile at a position (*l*_*x*_, *l*_*y*_) is given by:

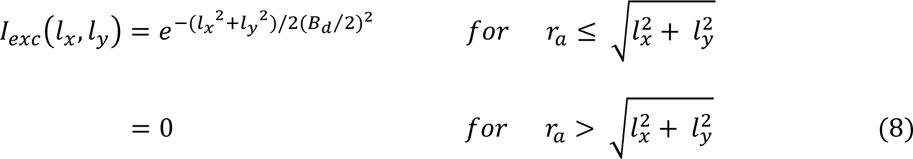

Furthermore, we assumed that the sources of autofluorescence are continuously and homogeneously distributed over the entire sample. A source of autofluorescence, located at position (*l*_*x*_, *l*_*y*_) in a plane parallel to the *xy*-plane at *z*, emits light when exposed to the excitation beam (this depends on the size of the excitation beam; see below) that propagates through the sample as a high-NA Gaussian beam (with focus at the location of the source), creating a Gaussian intensity profile at the focal plane (the *xy*-plane at *z* = 0), given by:

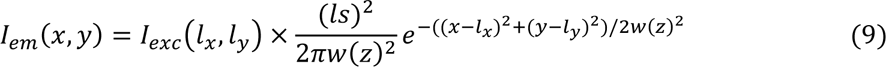

where, *ls* is the lattice spacing of sources (100 nm in our simulations) and *w*(*z*) is the width of the propagating beam *w*(*z*) at *z* = 0, given by equation (1). Here, the normalized intensity is multiplied by the intensity of the excitation beam at position (*l*_*x*_, *l*_*y*_). The total BG intensity contributed to the focal plane (*z* = 0) by a parallel plane at *z* is obtained by summing the intensity contributions from all individual sources located in the *xy*-plane at *z*. In this manner, we obtain the BG intensity contributed at the origin (*x*=*y*=*z*=0) by different out-of-focus *xy*-planes, moving along *z* (between 0 mm to 1 mm) with 100 nm steps. Summing the intensities at the origin contributed by all *xy*-planes, by taking the area under the curve, gives the total BG intensity at the origin, for a given excitation beam width. In the simulations, we scan through the height only in one direction, as we assume that the BG is the same from above or below the focal plane.

## Acknowledgements

We thank Dr. Andreas Biebricher for help with measuring the beam profile. We acknowledge financial support from the European Research Council under the European Union’s Horizon 2020 research and innovation programme (Grant agreement no. 788363; “HITSCIL”) and Marie Sklodowska-Curie Actions Postdoctoral Fellowship of the European Commission (Project no. 898006; ‘MingleIFT’, A.M.).

---

**Supplementary Figure 1:**
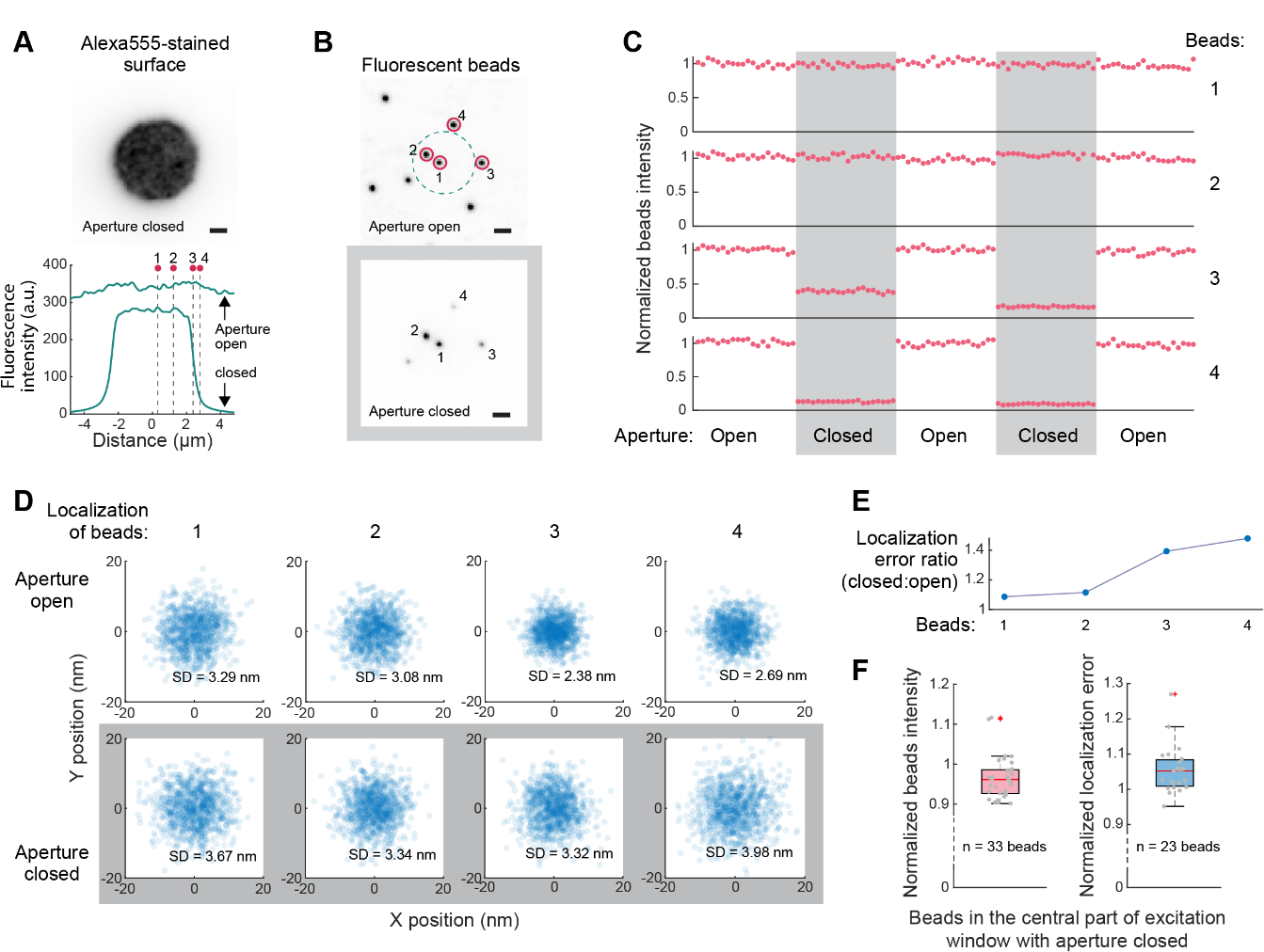
Effect of shape and size of the excitation beam on fluorescence signal intensity and localization precision. **(A)** Alexa555-coated surface excited with a cropped 491 nm beam of width ∼5 µm. Bottom: Alexa-555 fluorescence intensity measured with the aperture open and closed. The intensity is the central part is ∼1.3x lower when the aperture is closed. Red dots and dotted lines indicate approximate positions of fluorescent beads shown in B, with respect to the beam intensity profile. **(B)** Fluorescent microspheres imaged with aperture open (top; beam width ∼30 µm) and closed (right; excitation window width ∼4-5 µm). Dotted line approximately indicates the area illuminated when aperture is closed. Scale bar: 1 μm. **(C)** Normalized fluorescence intensity (BG subtracted) for the four beads indicated in A, imaged with aperture open and closed. For beads 1 and 2 located close to the centre of the beam, signal intensity does not change noticeably on changing aperture width. Signal intensity of beads 3 and 4 critically depends on the width of the aperture. Note: Slightly different aperture widths result in different intensity during the first and second “Closed” positions. **(D)** Localization of the four beads shown in A, tracked over 1000 frames, with aperture open (top) and closed (bottom). Localization error (SD) is indicated. **(E)** The ratio between localization errors with closed and open aperture shown in D. The ratio increases from ∼1.1 to ∼1.4 as beads get closer to the edge of the excitation window. **(F)** Fluorescence intensity (BG subtracted; left) and localization error (right) measured for beads located near the centre of the excitation window (such as beads 1 and 2 in A) with aperture closed, normalized to corresponding values with aperture open. There is ∼4% decrease in fluorescence intensity and ∼5% increase in localization error when the aperture is closed.

**Supplementary Figure 2:**
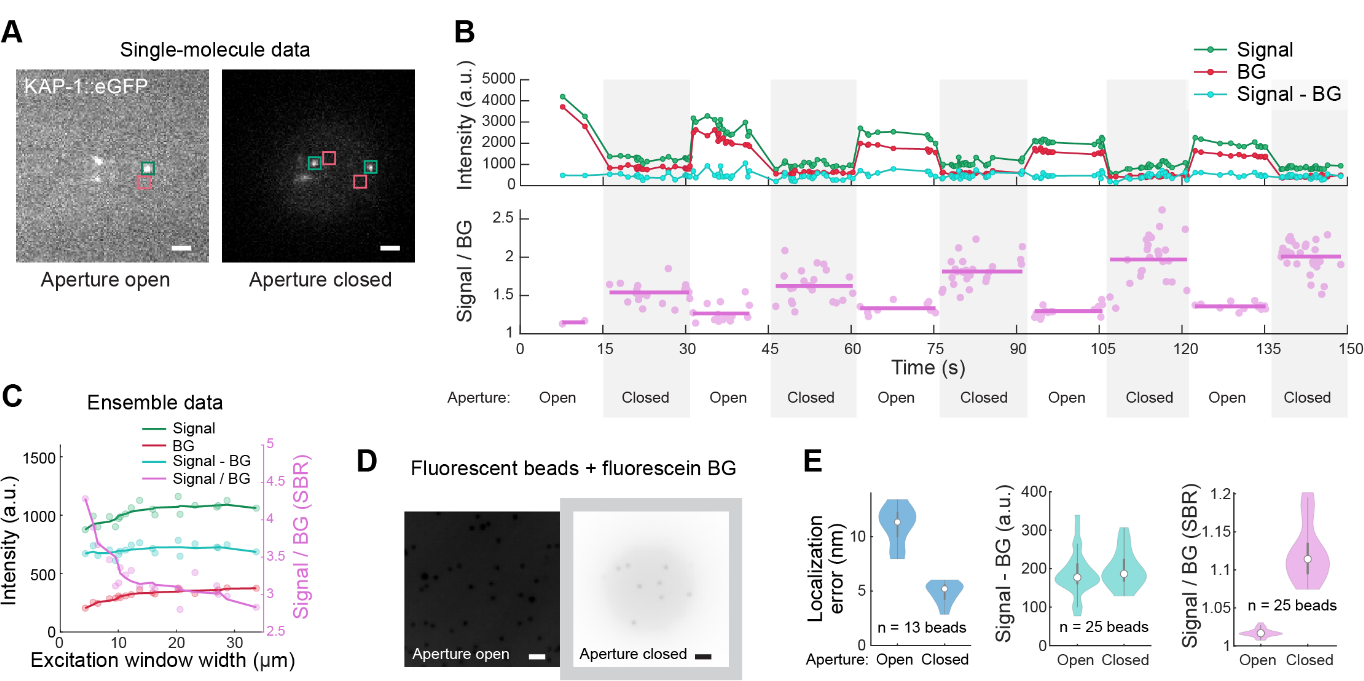
Effect of excitation window width on the signal and background intensities. **(A)** Example image frames from single-molecule imaging of KAP-1::eGFP with aperture open (left) and closed (right). Regions in which signal and BG intensities were measured are indicated in green and red, respectively. Scale bar: 1 μm. **(B)** Top: Signal, BG, and Signal-BG of single KAP-1::eGFP molecules as a function of time. Bottom: SBR measured for individual frames (circles) and the mean value at different aperture states (horizontal lines). The aperture state is indicated below. **(C)** Signal, BG, Signal - BG (left y-axis), and SBR (right y-axis) measured for KAP-1::eGFP, similar to Figure 2B-C, at different excitation window widths. **(D)** Same inverted images as shown in Figure 2F, but with brightness and contrast scaled the same. **(E)** Localization error (left), BG-corrected fluorescence intensity (Signal – BG; middle) and SBR (right) for several beads imaged, with fluorescein in the BG, with aperture open and closed. Aperture state is indicated below the x-axis.

**Supplementary Figure 3:**
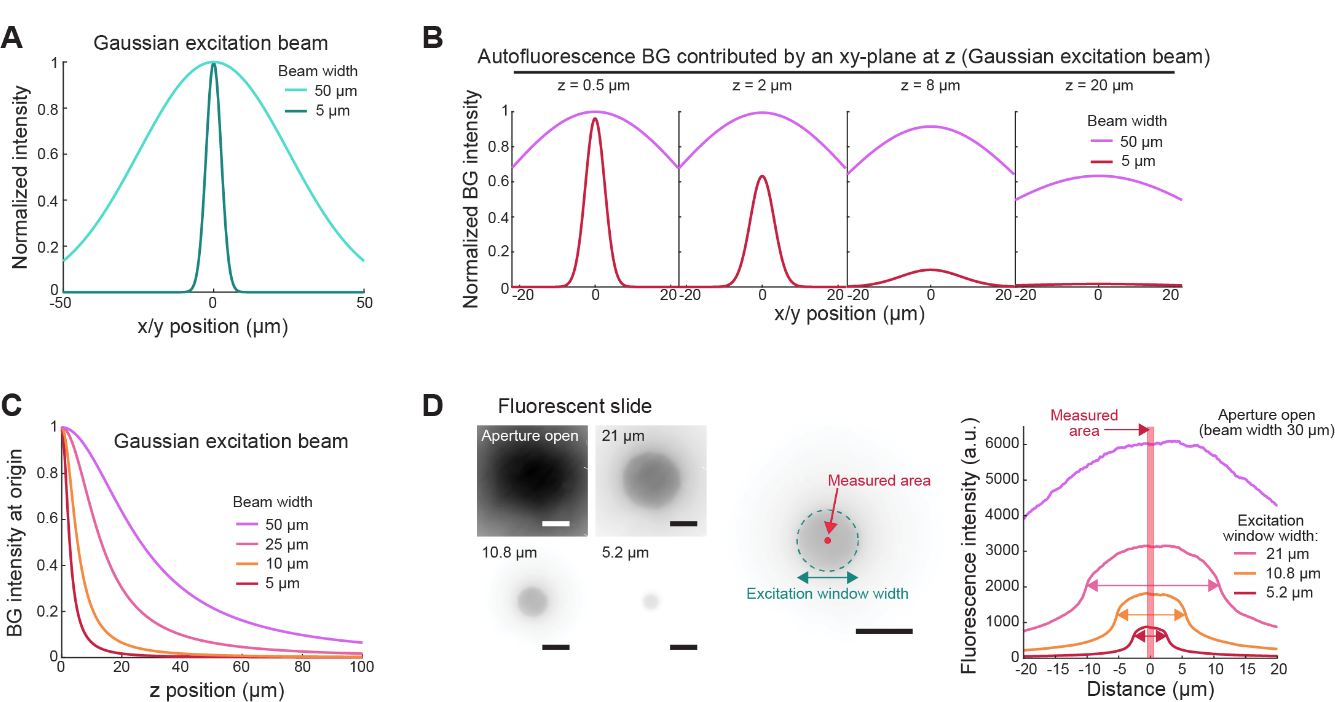
Details of the numerical simulations and experiments explaining the change in BG fluorescence with the width of the excitation beam. **(A)** The normalized intensity profiles of Gaussian excitation beams with width 50 μm (light cyan) and 5 μm (dark cyan). **(B)** BG intensity contributed by individual xy-planes (at z = 0.5 μm, 2 μm, 8 μm and 20 μm) illuminated by a Gaussian excitation beam with width 50 μm (magenta) or 5 μm (red). **(C)** Relative contribution to BG at the origin, from xy-planes at different heights (z) with respect to the focal plane (z=0). Sample illuminated by Gaussian excitation beams with indicated widths. **(D)** Left: Images of a ∼1.6 mm thick fluorescent slide illuminated by an excitation beam (width ∼30 μm) cropped by an aperture of varying width. Excitation window width is indicated; brightness and contrast are the same for all images. Middle: Excitation window width estimated as diameter of a circle drawn around the region with uniform high intensity. Intensity is measured in the small area in the centre of the illuminated region (red circle). Scale bar (left, middle): 10 μm. Right: Fluorescence intensity profiles for images shown on the left. Arrows indicate excitation window widths estimated as shown in the middle image. Light red rectangle indicates the width of the measured area shown in the middle image.

**Supplementary Figure 4:**
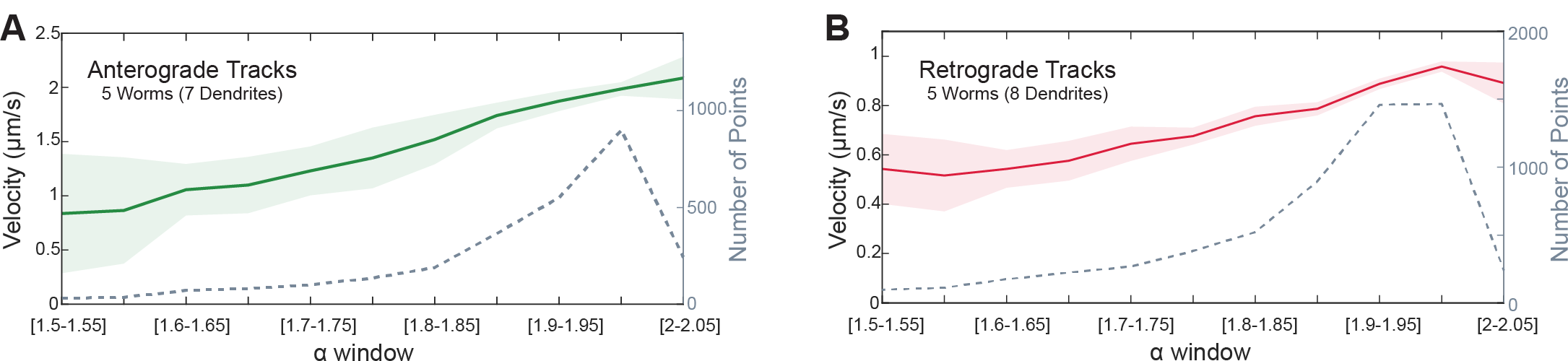
Velocity of anterograde and retrograde tracks of OCR-2-associated vesicles, filtered based on the local α-values associated with the tracks. **(A)** For anterograde tracks, average velocity of data points binned over alpha windows with bin width = 0.05, moving from 1.5 to 2.05, show that velocity increases from ∼1 μm/s to ∼2.1 μm/s. Number of datapoints in each bin is indicated by the dotted grey line (right y-axis). **(B)** For retrograde tracks, average velocity of data points binned over alpha windows with bin width = 0.05, moving from 1.5 to 2.05, show that velocity increases from ∼0.55 μm/s to ∼0.95 μm/s. Number of data points in each bin is indicated by the dotted grey line (right y-axis). For all distributions, average value and error are estimated using bootstrapping.

**Supplementary Figure 5:**
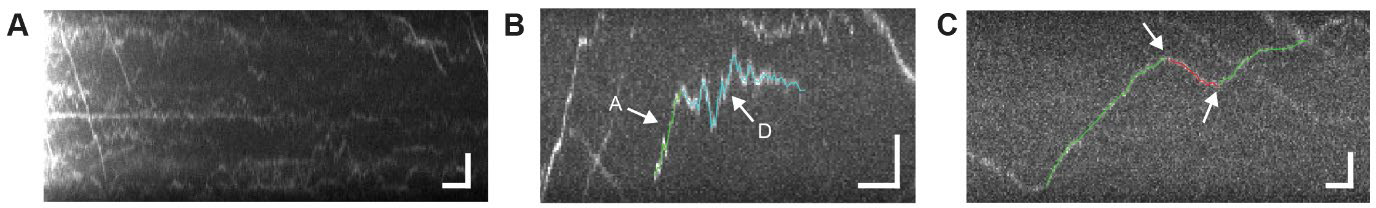
Diffusive tracks of OCR-2-associated vesicles. **(A)** Kymograph along the dendrite taken at the start of imaging showing the initial bleaching of many diffusive and stuck OCR-2-associated vesicles. Scale bars: 3 s (horizontal), 2 μm (vertical). **(B)** Kymograph, overlayed with track data, of a vesicle that first demonstrates directed motion in the anterograde direction along the dendrite (green, A), whereafter it switches to a diffusive state (cyan, D). Scale bars: 3 s (horizontal), 2 μm (vertical). **(C)** Kymograph, overlayed with track data, of a vesicle that shows directional switching from anterograde (green) to retrograde (red) and back to anterograde (green) as indicated by the arrows. Scale bars: 1 s (horizontal), 2 μm (vertical).

**Supplementary Movie 1: Imaging OCR-2-associated vesicles in the dendrite of a sensory neuron in *C. elegans*, using SWIM and epifluorescence microscopy**. OCR-2::eGFP dynamics in the dendrite were imaged with a closed aperture (excitation window width ∼7 µm) for the first 15 min. After 15 min, the aperture is opened (beam width ∼30 µm), with imaging performed for another 5 min. Four sections of the acquired stream (acquisition rate 6.67 fps) are shown, with time indicated in min:sec (5x sped up). Brightness and contrast of section 3 are different from the other fragments.

**Supplementary Movie 2: OCR-2-associated vesicles imaged in the dendrites of sensory neurons in *C. elegans*, using SWIM**. OCR-2::eGFP dynamics in a pair of PHA/PHB dendrites (labelled Dendrite 1, Dendrite 2) imaged with the aperture closed (excitation window width ∼11 µm) for 15 min at frame rate 6.67 fps. Time is indicated in min:sec (7.5x sped up).

**Table S1:**
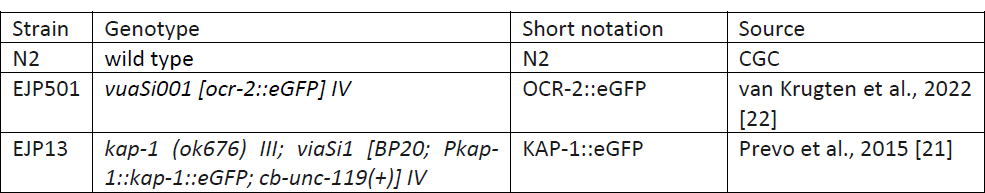
*C. elegans* strains used in this study.

## Notes

### Competing Interest Statement

The authors have declared no competing interest.

### Summary of Updates

In the revision we have characterized our microscopy technique more rigorouly, performing several additional controls. Furthermore, we have added a completely new section in which we show theoretical calculations that explain the significant reduction in background fluorescence on reducing the window-width of the illumination window.

